# Foldclass and Merizo-search: embedding-based deep learning tools for protein domain segmentation, fold recognition and comparison

**DOI:** 10.1101/2024.03.25.586696

**Authors:** S. M. Kandathil, A. M. Lau, D. W. A. Buchan, D. T. Jones

**Author notes:** **Corresponding Author**: David T. Jones. InstaDeep Ltd, 5 Merchant Square, London, W2 1AY, UK.

## Abstract

The availability of very large numbers of protein structures from accurate computational methods poses new challenges in storing, searching and detecting relationships between these structures. In particular, the new-found abundance of multi-domain structures in the AlphaFold structure database introduces challenges for traditional structure comparison methods. We address these challenges using a fast, embedding-based structure comparison method called Foldclass which detects structural similarity between protein domains. We demonstrate the accuracy of Foldclass embeddings for homology detection. In combination with a recently developed deep learning-based automatic domain segmentation tool Merizo, we develop Merizo-search, which first segments multi-domain query structures into domains, and then searches a Foldclass embedding database to determine the top matches for each constituent domain. Combining the ability of Merizo to accurately segment complete chains into domains, and Foldclass to embed and detect similar domains, Merizo-search can be used to detect per-domain similarities for complete chains. We anticipate that these tools will enable a number of analyses using the wealth of predicted structural data now available. Foldclass and Merizo-search are available at https://github.com/psipred/merizo_search. Merizo-search is also available on the PSIPRED web server at http://bioinf.cs.ucl.ac.uk/psipred.

## Introduction

In the post-AlphaFold2 (Jumper *et al*., 2021) era, the field of structural biology has been revolutionised by an influx of predicted structures, generated at the push of a button. Consider this question: given a protein of interest, what is the best way to identify similar structures from within a database of more than 200 million AlphaFold2, or 600 million ESMFold (Lin *et al*., 2023) structures? In such a situation, most, if not all traditional methods for structural comparison (e.g. TMalign (Zhang and Skolnick, 2005) and SSAP (Orengo and Taylor, 1996)), will struggle with the sheer volume of data. This challenge is partially mitigated by advanced tools like Foldseek (van Kempen *et al*., 2023), which navigates the extensive search space far more rapidly than traditional tools. Foldseek’s efficiency stems from its unique encoding system, which compresses the contact information of a protein’s 3D structure into a one-dimensional sequence (or “3Di string”), enabling it to make use of highly efficient alignment and clustering methodologies introduced in its predecessor MMseqs2 (Steinegger and Söding, 2017). However, fast as it is, Foldseek is not without its limitations. Its simplified representation of 3D structures, relying on nearest neighbour contacts, may inadvertently lead to similar 3Di strings generated for certain unique but similar protein folds.

This consideration is particularly relevant in the context of structural homology detection, which concerns the detection of distant evolutionary relationships between pairs of proteins, seemingly unrelated by sequence, but which are related by the fold of their structures. Many methods have been devised to aid in this task, leveraging state-of-the-art deep learning methodologies for comparing structures and include methods such as TMVec (Hamamsy *et al*., 2023) and Progres (Greener and Jamali, 2022). In short, these methods each make use of embedding-based approaches for determining similarity within an embedding space. A neural network embeds the protein sequence, structure, or both, into a latent embedding space, whereby similar proteins are localised to the same areas of the embedding space during training. Distance metrics such as the cosine distance can be used to calculate a single score representing the similarity between two embeddings in a latent space (i.e. two structures). A similar but machine learning-free approach is taken in Geometricus (Durairaj *et al*., 2020), which decomposes a query structure into a set of fragments and 3D moment invariants that characterise local structural neighbourhoods. These fragments are discretized, and using the set union of such fragments observed in a reference set of proteins, and the vector of counts of each fragment serves as an embedding of the structure, which can then be searched against a precomputed database of such embeddings to find similar structures.

However, the task of structure searching is further complicated by a multitude of factors. Many proteins consist of multiple domains that can appear in varying orders and conformations, necessitating a clear definition of “similarity” between two structures. To address this challenge, searches can be conducted at the domain level by initially decomposing a query protein into its constituent domains before comparing them against a library of annotated domains, such as those found in CATH (Sillitoe *et al*., 2021) and ECOD (Cheng *et al*., 2014). As domains constitute the functional units of proteins, this approach further facilitates the transfer of functional annotations by linking specific regions of a protein to known functional domains. Automated domain parsing methods like Merizo (Lau *et al*., 2023), Chainsaw (Wells *et al*., 2023), UniDoc (Zhu *et al*., 2023), and SWORD (Postic *et al*., 2017) can be employed to accomplish this task, with the newer methods Merizo and Chainsaw capable of operating on both experimental structures and those predicted by deep learning methods such as AlphaFold. This distinction is crucial, as models generated by methods such as AlphaFold2 and ESMFold may feature long stretches of unstructured regions that are not part of domains (non-domain residues; NDR), owing to the full sequence being modelled. With few exceptions, NDRs are typically absent in experimentally determined structures.

We describe a new method for conducting protein structure similarity searching at the domain level. This tool extends the functionality of our recent method Merizo, allowing a multidomain query to be segmented using Merizo before embedding domain structures into a latent representation using an equivariant graph neural network called Foldclass. Domain embeddings are compared against a library of precomputed structure embeddings from structure databases such as the CATH database, and the top *k* nearest neighbours (measured by cosine distance) are returned, with further validation using TMalign to confirm hits. Since Merizo has already been benchmarked for accuracy on domain segmentation tasks (Lau *et al*., 2023), here we focus on the development and performance of Foldclass for embedding-based similarity searching, and its integration with Merizo into Merizo-search.

## Results

### Integration of Merizo and Foldclass in Merizo-search

A summary of our combined Merizo and Foldclass method is shown in Figure 1. Our method consists of several key routines which make it convenient for a multidomain query to be searched against a library of domains. Firstly, given a multidomain query, the Merizo-segment routine can be called to predict the domains of the query. These domains are then encoded using the Foldclass network into fixed size embedding of shape [N, 128], where N is the number of identified domains in the original query. Query domain embeddings are then searched against a structure library which can be separately prepared using the Merizo-createdb routine, which handles the high-throughput embedding of each structure into a custom Foldclass database containing the per-structure embeddings of size [M, 128] (where M is the number of encoded target structures), and their Cα coordinates. The Merizo-search functionality conducts the searching of the query domain embeddings against the target embeddings and the distance between all queries and targets are computed via a metric such as the cosine distance. In the final step, Merizo-search validates the top *k* targets (ranked via the distance metric in the previous step) by computing the TM-align score between their Cα coordinates. To make our method more comprehensive, the Merizo easy-search method can be used to automatically run Merizo-segment and Merizo-search on a query structure, provided the user already has a pre-encoded target database.

**Figure 1.**
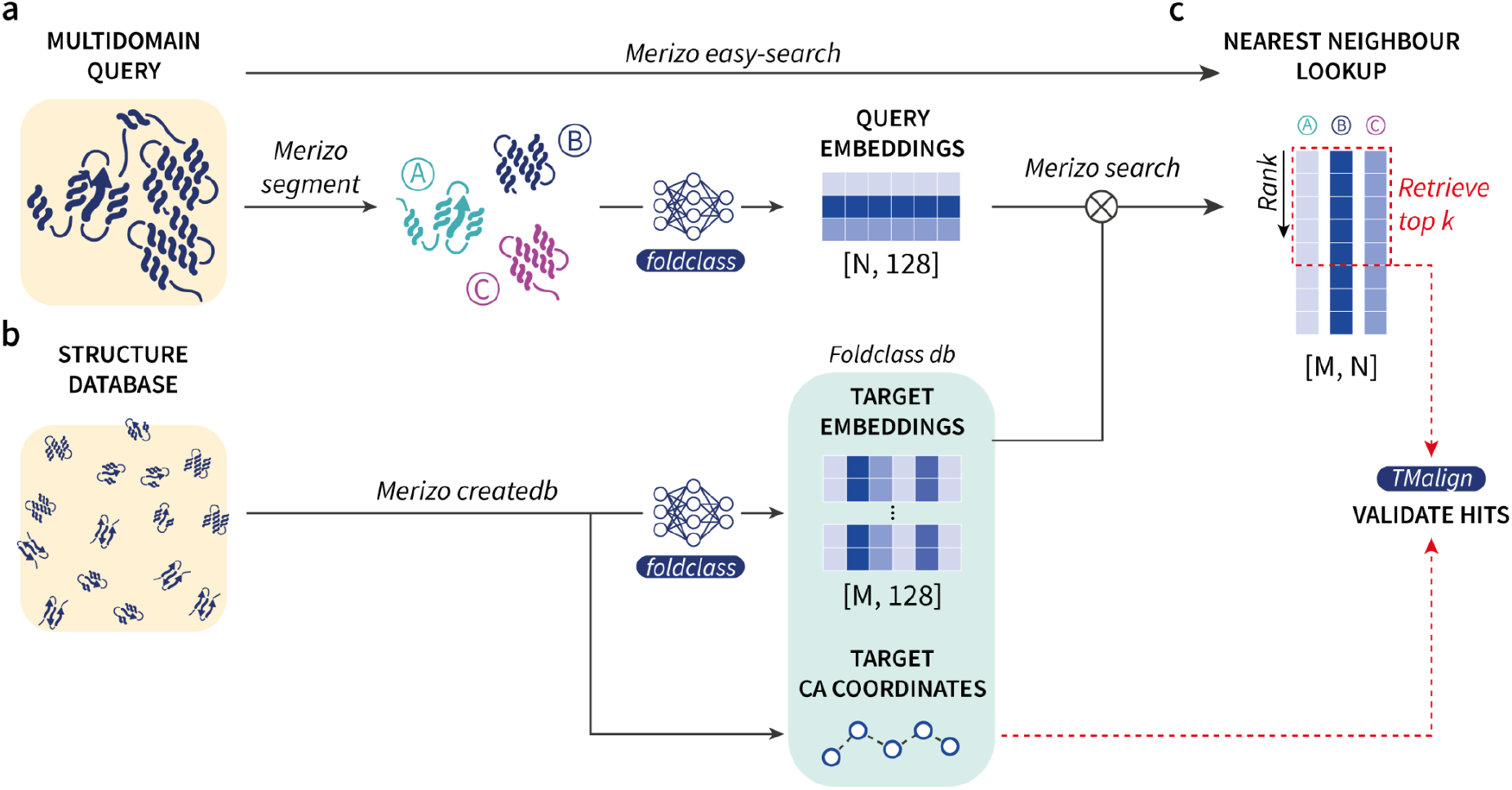
Workflow of Merizo-search. (**a**) A query composed of multiple domains can be input to the *segment* module of Merizo-search which segments the query into *N* domains using Merizo. Domains are then encoded using Foldclass into fixed-size embeddings. (**b**) The *createdb* module allows a structure database (e.g. CATH domains) to be encoded into a database of embeddings and Cα coordinates using Foldclass. (**c**) The *search* module computes the nearest neighbour of each query embedding against target embeddings using cosine distance. The nearest neighbours of each query domain is ranked according to cosine distance and the top *k* results are validated using TM-align. The *easy-search* module uses the *segment* and *search* routines to directly search a multidomain query against a pre-encoded target database of structures.

### Accuracy of Foldclass embeddings for structure similarity searching

To validate the Foldclass embedding network itself, we looked at its classification performance on a holdout set of 62 domains from the last 3 CASP experiments for which no homologs could be found in CATH, but which did have matches at the fold (T) level. On this set, Foldclass has an overall accuracy of 45/62 (73%). However, when thresholded at a confidence of 0.9, 42/44 (precision 95%) of the domains are correctly labelled (recall 65%), showing that the Foldclass network performs well on these hard cases when used as just a simple classifier.

We then compared the performance of Merizo-search against Foldseek, in both TMalign and 3Di+AA alignment modes. In each experiment, we search the CATH (v4.3) S40 representative domains against the CATH S20 set and evaluate whether each S40 query is matched with a S20 target of the same CATH superfamily. As not all CATH superfamilies are included within the CATH S20 and S40 representative sets, we evaluate only domains belonging to superfamilies which are present in both datasets (n=21,005). For each S40 query, we take its corresponding CATH superfamily label and check the top target returned by each method (sorted by descending query TM-align score). A match is considered a hit if the target domain belongs to the same CATH superfamily.

The results shown in Figure 2 indicate that Merizo-search identifies a comparable number of hits to Foldseek when the value of top *k* is increased, with the most hits identified at *k*=20. Compared to Foldseek, Merizo-search outperforms both the ‘fast’ and ‘fastest’ search modes of Foldseek which perform faster searches but at great cost to sensitivity.

**Figure 2.**
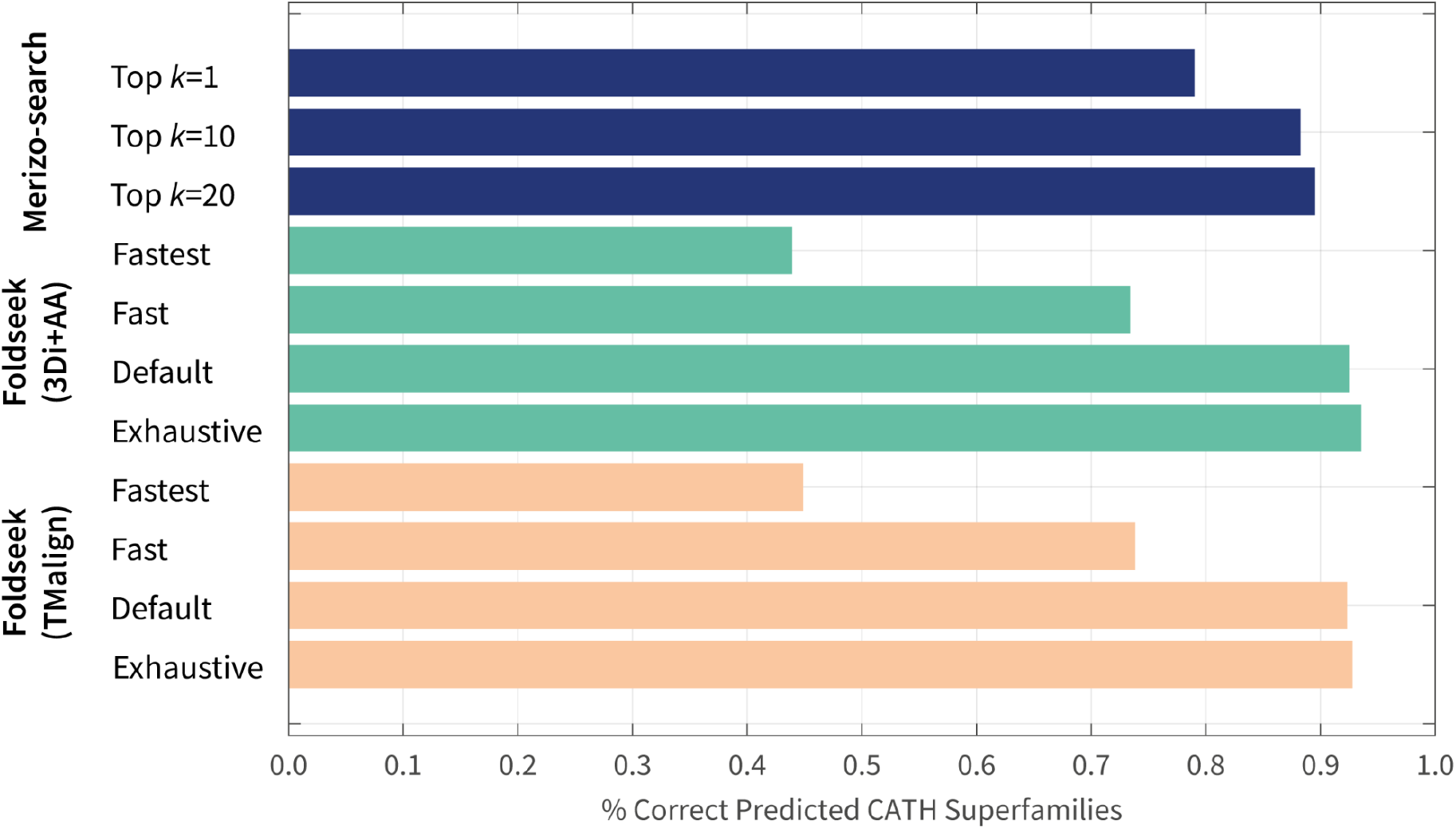
Comparing Merizo-search and Foldseek on CATH superfamily predictions. Each bar represents the percentage of queries where the top ranked target belongs to the same CATH superfamily (n=21,005).

We further evaluated whether the hits identified by Foldseek and Merizo-search overlapped. The results of this benchmark are shown in Tables 1 and 2 for Foldseek in TM-align modes and 3Di+AA modes respectively. Overall, we see that Merizo-search is able to recover several hundred to several thousand queries that Foldseek cannot find correct matches for, with the highest recovery gained when Foldseek is used in its “fast” and “fastest” modes (which differ from the default mode by altering the sensitivity setting). Compared to Foldseek’s exhaustive modes, Merizo-search is able to find matches for several hundred more queries, even when running Merizo-search in the least sensitive *k*=1 mode. Taken together, these results suggest that structure matching via embedding similarity can be used to complement Foldseek in order to maximise coverage for a set of queries.

**Table 1.** Number of additional hits recovered by Merizo-search over Foldseek (TMalign mode). n=21,005.

|  | Merizo-search |  |  |
| --- | --- | --- | --- |
| Foldseek | Top k=1 | Top k=10 | Top k=20 |
| Fastest (s=1) | 8140 | 9134 | 9285 |
| Fast (s=4) | 3004 | 3600 | 3707 |
| Default (s=9.5) | 147 | 191 | 211 |
| Exhaustive | 111 | 134 | 151 |

**Table 2.** Number of additional hits recovered by Merizo-search over Foldseek (3Di+AA mode). n=21,005.

|  | Merizo-search |  |  |
| --- | --- | --- | --- |
| Foldseek | Top k=1 | Top k=10 | Top k=20 |
| Fastest (s=1) | 8948 | 10002 | 10158 |
| Fast (s=4) | 3313 | 3994 | 4110 |
| Default (s=9.5) | 240 | 378 | 414 |
| Exhaustive | 124 | 227 | 255 |

### Evaluation of multi-domain searches

Domain-centric embedding and searching potentially allows us to carry out multi-domain structure search while being robust to rearrangements of the individual domains in different chain structures. We implemented a simple means to find database chains that contain all the domains in a segmented query chain, or that contain all of a set of user-specified domains, and evaluated its performance in terms of precision of finding hits with the correct multi-domain architecture in the list of hits for each multi-domain query structure (Methods).

Table 3 compares the mean precision for the multi-domain hits identified by Merizo-search and Foldseek, comparing a few different settings of each method (Methods). We find that Merizo-search achieves the highest precision for recognising multi-domain hits, especially when only hits reported as being exact multi-domain architecture (MDA) hits are considered. Precision drops slightly when the value of *k* is increased, or when considering queries with more than 2 domains, reflecting an increased false-positive rate due to the inexact nature of the embedding-based search. For both Foldseek and Merizo-search, precision is also higher for T-level matches as compared to H-level matches, reflecting the fact that H-level matches require more intensive searches involving structure and sequence to accurately identify. Nevertheless, these results show that Merizo-search achieves precise identification of multi-domain hits at fold-or superfamily-level. Notably, this level of precision is achieved following automatic segmentation by Merizo, indicating that the overall search procedure is fairly robust to occasionally inaccurate domain parsing.

**Table 3.** Mean precision when retrieving multi-domain hits at either superfamily (H) or fold (T) level. Results are shown for all query chains (first two columns) and for query chains with more than 2 domains (n_dom_>2; last two columns). The highest value in each column is shown in bold.

|  |  | Mean precision |  |  |  |
| --- | --- | --- | --- | --- | --- |
| | | H-level | T-level | H-level,<br>$n_{\text{dom}} > 2$ | T-level,<br>$n_{\text{dom}} > 2$ |
| Foldseek | k=1000 (default) | 0.609 | 0.649 | 0.581 | 0.600 |
|  | k=100 | 0.729 | 0.758 | 0.696 | 0.713 |
|  | k=500 | 0.637 | 0.674 | 0.605 | 0.624 |
| Merizo-search<br>(easy-search<br>mode) | k=100, any match | 0.776 | 0.816 | 0.785 | 0.808 |
|  | k=500, any match | 0.703 | 0.756 | 0.740 | 0.768 |
|  | k=100, MDA match | **0.865** | **0.910** | **0.831** | **0.850** |
|  | k=500, MDA match | 0.821 | 0.887 | 0.800 | 0.826 |

The multi-domain search procedure of Merizo-search categorises hits according to the relative sequential arrangement of the domains in the query and hit chain (Methods), the most general being a “bag-of-domains” hit, in which all query domains are matched non-sequentially to those in the hit chain, which may also have extra, unmatched domains. In general, Merizo-search reports per-domain correspondences between the query and hit independent of the order of the domains in each chain, allowing users to inspect the hit list further to identify match patterns other than just exact MDA matches. An example of such a non-sequential multi-domain match is shown in Figure 3 for the query chain 3bmvA (Cyclomaltodextrin glucanotransferase from *B. subtilis*) and the hit chain 2e8zA (Pullulanase from *B. subtilis* strain 168). All four domains in the query chain 3bmvA match the domains in the hit chain 2e8zA with a TM-align score exceeding 0.5, and their CATH annotations match up to the fold level. The order of the corresponding domains is different between the two proteins, and the three-dimensional organisation of the domains because of this, Foldseek is unable to identify the chain 2e8zA as a hit. Both chains are carbohydrate-degrading enzymes, suggesting a possible evolutionary link. This illustrative example shows that the domain-centric matching strategy in Merizo-search can be used as a starting point for analyses of domain relatedness and evolution in different multi-domain contexts.

**Figure 3.**
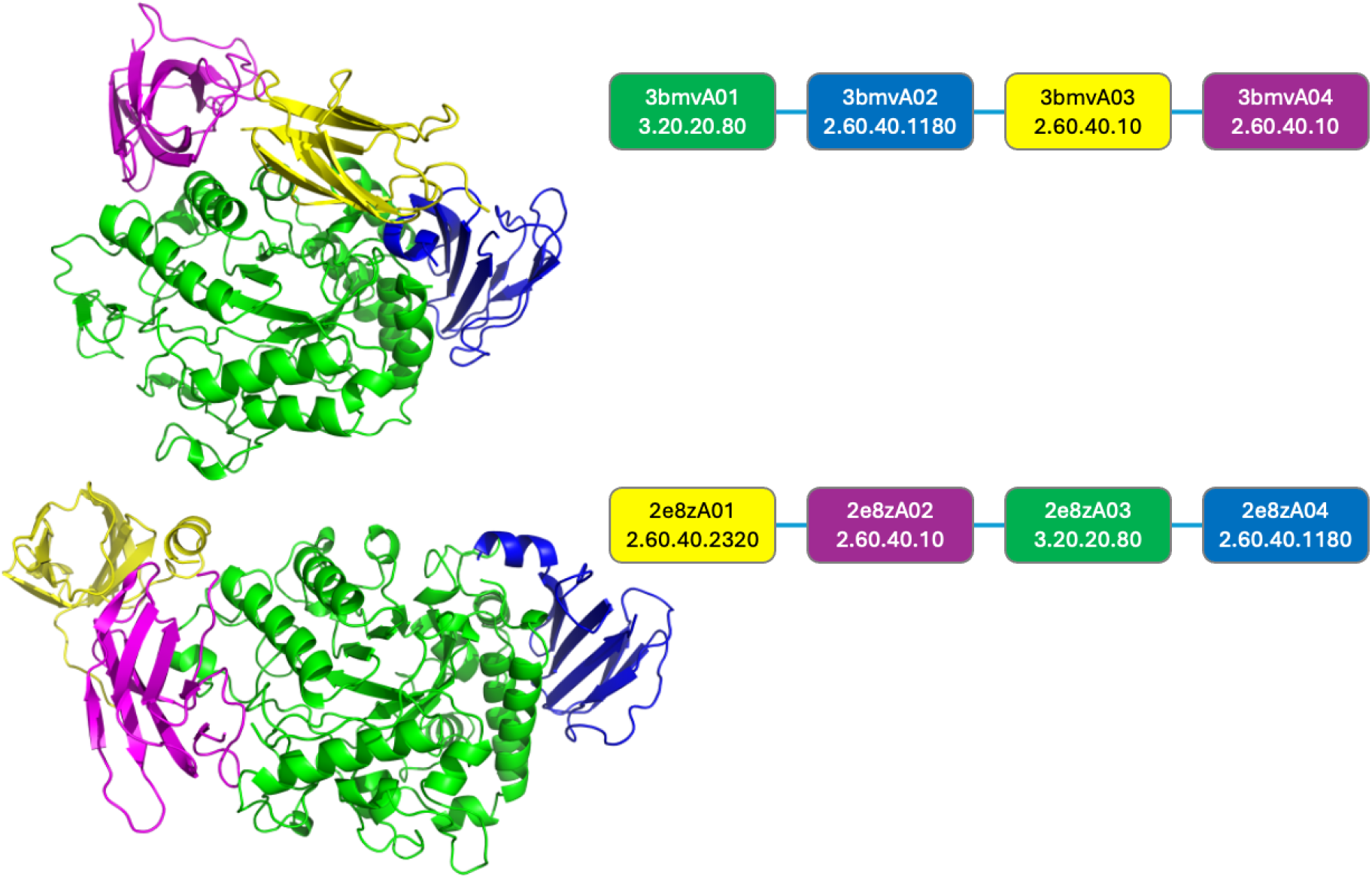
Example of a “bag-of-domains” match retrieved by the multi-domain search mode of Merizo-search. The query chain (3bmvA, top) and a hit chain (2e8zA, bottom) are shown, together with a schematic representation of the multi-domain architecture in each chain sequence, including CATH domain annotations. Colours in the structure cartoons and schematics indicate domain correspondences identified by Merizo-search.

## Discussion

The ability to represent protein domains as fixed-length vectors lends itself to a number of applications, including the ability to rapidly conduct similarity searches against libraries of domains. To date, most embedding methods for protein sequences and structures have focused on per-residue embeddings computed for entire chains, meaning that comparison is most accurate and meaningful when comparing entire proteins. A more complete picture of evolutionary relatedness would need to take into account the domain architecture of the proteins being compared. Our work using Foldclass illustrates the principle that this technique for searching is indeed effective, and the strategy we employ lends itself to more scalable approaches that can meet the challenge of searching very large databases of protein domain structures. The ability to do this opens up new possibilities in furthering our understanding of the protein universe. Additionally, we leveraged our recently developed domain segmentation tool, Merizo, to first automatically segment a query structure if needed, embed the constituent domains using the Foldclass neural network and then search the constituent domains against a precomputed database of Foldclass-derived embeddings. This integrated tool makes it convenient for users to assess domain-level evolutionary relationships between protein structures, and we anticipate that it can be used to discover similarities and differences between multi-domain structures, which account for a significant fraction of proteomes across the Tree of Life. Consideration of multi-domain architectures can be used to classify and detect relationships between proteins; for example, a recent study provides functional groupings of kinases by explicitly considering multi-domain architecture of the relevant proteins (Adeyelu *et al*., 2023). Such efforts can be accelerated and expanded using the methods developed in this work.

Merizo-search jobs can also be run on the PSIPRED web server (Buchan *et al*., 2024;) (http://bioinf.cs.ucl.ac.uk/psipred) in the easy-search mode. The web server shows a graphical representation of the domain segmentation produced by Merizo, as well as the top hits for each domain in the chosen domain database. Currently, the web server allows users to search all domains in CATH 4.3 (540,417 domains) or TED100 (∼324 million domains) (Lau *et al*., 2024).

## Methods

### Model architecture and Training

The Foldclass neural network model is essentially a stack of two or more E(n)-equivariant graph neural network (EGNN) blocks (Satorras *et al*., 2021), with the residue positional encoding used as the node features, and the original domain Cα coordinates used as the coordinate inputs. The EGNN layers compute updated node representations using a message-passing framework and have the property of being equivariant to rotations, translations, reflections and permutations of the input coordinates. In our implementation, we omit the coordinate update step used in the original, as this is not useful for a fold classification task, and the same unchanged coordinates are presented to all the EGNN blocks. The EGNN blocks use a hidden dimension of 128 and an output dimension of 256 for the node and edge update sub-networks. The updated node features from the final EGNN layer are then averaged to yield the final embedding of the input structure. This yields the final fixed-size embedding for a given input structure regardless of the number of residues.

Input node features are just sinusoidal positional encoding vectors (Vaswani *et al*., 2017) for the sequence. An encoding of the sequence itself was not used as input to the model, as this was found to cause rapid overfitting.

Unlike previous methods, which tend to make use of contrastive loss, Foldclass is initially trained purely as a multiclass classifier. This turns out to be a very efficient and stable way of training the model. The final average embedding of the input structure is used as the input for three separate linear layers. These three output heads of the network predict logits for the Class (C), Architecture (A) and Topology (T) level classifications in CATH. The classification heads are trained using categorical cross-entropy loss functions with inbuilt softmax functions. During training, class weights were set to f_min_/f_i_ i.e. weighted according to reciprocal frequency of each class in the training set where the minimum class frequency in the training set is given a weight of 1. Training the network on the different levels of the CATH hierarchy in parallel, where the C and C.A labels are effectively acting as auxiliary losses, reduces overfitting and gives a more robust final embedding.

The training set comprised 31,885 domains, which are the S30 non-redundant subset of domains in the CATH 4.3 release. A small random validation set of 50 was used to monitor overfitting. Although this classifier training is considered as pre-training of the model, we did make use of a separate classification test set of 62 domains that were targets in CASP13-15 which have no detectable homologs in the 4.3 release of CATH, but which did have matching folds and so could be assigned CAT labels.

The number of different labels for the 3 output heads are as follows: 5 (C), 43 (CA) and 1421 (CAT). The EGNN weights are initialised from a normal distribution with a mean of 0 and a standard deviation of 1e-3, while the classification layers are initialised using the PyTorch default initialization. The Foldclass network is trained for a maximum of 300 epochs using the AdamW optimiser (Loshchilov and Hutter, 2017; Kingma and Ba, 2014) with a learning rate of 0.0003 and a weight decay of 1E-2. Further regularisation is provided by adding Gaussian noise with a standard deviation of 1.5Å to the Cα input coordinates of each training example. No noise is added for validation samples.

### Using Foldclass for structure embedding

Although Foldclass is trained as a classifier, in practice its primary use is in generating an embedding vector, which is a representation of the overall chain fold of the protein domain. This is achieved by simply removing the linear layers which output the classification logits and taking the mean output from the EGNN blocks as the embedding vector. These vectors can easily be compared by say Euclidean distance or cosine similarity, like any embedding vector.

### Database construction and searching

A Foldclass database is a collection of the sequences of the constituent domain structures, along with their embeddings computed from the Foldclass neural network. The embedding of the query domain is then used to compute cosine similarities to the embeddings stored in the database. The top *k* hits with highest similarity are returned. The parameter *k* can be varied and we investigate the performance of a few settings of *k*. This approach is acceptable as a proof of principle, but does not scale to very large database sizes (>10M entries), for which we employ the Faiss library (Douze *et al*., 2024) for efficient searching. We found that exhaustive inner-product searches on normalised Foldclass vectors could be performed in acceptable time frames even when searching ∼365 million domains in the complete TED database (Lau *et al*., 2024) (roughly 2 min when the database is stored on fast NVMe storage and a GPU is used for searching). Therefore, we opted to use exhaustive inner-product searches for maximum accuracy when using databases too big to fit in system memory, and this retains consistency with the default PyTorch-based inner-product search used when Faiss is not used.

### Multi-domain searches

In addition to per-domain searches, we implemented a mechanism to find database hits that cover all query domains. When a set of domain structures is provided as a query, the database is first searched on a per-domain basis as above. We then retrieve all domains from the chains in the database that contained a hit to any query domain in the top-*k* lists (*k* still controls the maximum number of per-domain hits retrieved), discarding domains from chains in the database that have fewer domains than there are in the query chain/set. In this way, the initial per-domain embedding-based search serves as a prefilter to reduce the size of the set of database domains that can then be searched more intensively. We perform all-against-all TM-align comparisons between query and potential hit domains to determine possible matches in the reduced search space. Finally, we identify those chains in the database that match all domains in the query at a specified TM-align score threshold (0.5 by default), while allowing each domain in a database chain to be matched to exactly one query domain at a time. During this process, we enumerate all pairings of query and hit domains that meet the per-domain TM-align threshold, and classify multi-domain hits into one of four categories, each a subset of the last:

1. “Bag-of-domains” match: domains in the hit chain match all the query domains, but in any order and additional domains may be present in the hit chain.
2. Gapped domain alignment, type 1: domains in the hit chain match all the query domains in the same order as the query, but possibly with extra domains either at the chain termini, or in between the matching domains.
3. Gapped domain alignment, type 2: domains in the hit chain match all the query domains in the same order as the query, and any extra domains in the hit chain only occur at one or both termini.
4. Exact multi-domain architecture (MDA) match: The hit chain has the exact domain composition and order as the query chain, with no extra domains.

In the case of the ‘easy-search’ workflow, the complete multi-domain search procedure is carried out separately for each query chain after segmentation by Merizo. In such cases, accurate categorisation of the multi-domain hits into the above categories depends on the accuracy of the segmentation, as it depends on the number of domains identified in the query chain. In the case where a user supplies pre-segmented domain structures for searching, these are treated as originating from a single chain in the order that they are provided.

We evaluated the ability of the multi-domain search to retrieve hits from a set of 312,544 PDB chains in CATH 4.3 which have no segmented but unclassified domains, and for which full-chain structures could be retrieved. The set contains both single- and multi-domain chains. Using a set of 1828 multi-domain chains of distinct multi-domain architectures as queries, we tested the ability of the ‘easy-search’ mode of Merizo-search to segment each query chain and subsequently identify chains in CATH containing matches for all identified domains, at both the H-level and the T-level, considering any hit or only exact MDA matches (category 4 above). Since Merizo-search lists alternative query-hit domain pairings between the same query and hit chain (e.g. in the case of chains with tandem repeat domains), we only count a single hit chain per query chain. A hit chain was considered a true positive if its CATH annotations at the relevant level (either T or H) matched the ground-truth CATH annotations for the query chain. Results were reported for all the query chains, and considering only query chains with more than two domains (n = 1159).

We compared our results against Foldseek, using a Foldseek-formatted database of the same set of 312,544 CATH chains as used for constructing the Merizo-search database. Note that the Foldseek database used for this experiment comprises complete PDB chains, whereas the Merizo-search database contains domains, as Foldclass embeddings are trained on domains. For Foldseek, we used the 3di+AA alignment type on full-length chains, using the following command line:

~~~
foldseek easy-search query_chains/ cath_multidomain_chains_db
output_${k}.m8
tmp_foldseek --max-seqs $k -c 0.8 --cov-mode 2 --alignment-type 2 –e
0.01 --format-output
query,target,qlen,tlen,qcov,tcov,qtmscore,ttmscore,alntmscore,rmsd,e
value,bits
~~~

Where the setting of --max-seqs (‘$k’) is set to 100, 500, or the default of 1000. This parameter controls the maximum number of database chains per query that are allowed to pass the Foldseek pre-filter. Merizo-search uses per-domain searches as a pre-filter to reduce the initial search space, so the setting of *k* for Merizo-search is loosely analogous to that used for Foldseek, though they are not exactly equivalent. For completeness, we also evaluate Foldseek using the default setting of the --max-seqs parameter.

## Notes

https://github.com/psipred/merizo_search

### Competing Interest Statement

The authors have declared no competing interest.

### Summary of Updates

New functionality in the Merizo-search program has been described and evaluated (multi-domain search mode). Information on web server availability has also been added.

